# Real-life ear-EEG Recordings of Auditory Responses to Ambient Sounds

**DOI:** 10.64898/2025.12.02.691416

**Authors:** Jacopo Comoglio, Bowen Duan, Astrid Rands Bertelsen, Simon Lind Kappel, Preben Kidmose

**Affiliations:** Department of Electrical and Computer Engineering, Aarhus University, Finlandsgade 22, 8200 Aarhus N, Denmark

## Abstract

Despite being one of the most researched neural responses, auditory potentials have primarily been measured in controlled laboratory environments. However, there is a big interest in measuring auditory potentials in more naturalistic environments, with stimuli resembling, to a higher degree, natural sounds.

The objective of this study is to investigate auditory evoked potentials elicited by naturally occurring sounds in real-life environments. The paper presents an integration of a portable hearing aid research platform and a portable ear-EEG platform. This setup was validated through recordings of an Auditory Steady State Response (ASSR), elicited by real-time amplitude modulation of the ambient sound at 40Hz. Recordings were conducted in real-life under three conditions: walking and sitting with open and closed eyes.

In addition to ear-EEG, scalp EEG was recorded as a reference. For analysis, EEG signals were categorized into three groups: scalp channels, cross-ear channels, and within-ear channels. For each condition and channel group, the signal-to-noise (SNR) of the ASSR was calculated. The ASSR was statistically significant across all channel groups and conditions. A post-hoc analysis assessed the recording duration required to obtain a significant ASSR, revealing that 10 minutes were enough for within ear ear-EEG for all but two cases, whereas less than 7 minutes were sufficient for all but one case when using scalp-EEG.

**Clinical relevance:** The Auditory Steady-State Response (ASSR) is a versatile tool for assessing various characteristics of the auditory pathway. The ability to measure ASSR in real-life conditions using ambient sound could facilitate the development of hearing aids capable of dynamically adapting to the user’s current hearing loss.

## I. INTRODUCTION

Auditory Evoked Potentials (AEPs) are one of the most well researched brain responses [1][2]. However, most research on AEPs have been performed in laboratory settings, where the participant is often asked to stay still and presented with unnatural stimuli. One of the limiting factors for the use of natural stimuli, is the lack of a specific instrumentation to present highly controlled auditory stimuli in a mobile setup, resulting in scientists having to develop their own custom portable hardware [3][4][5]. Additionally, using natural ambient sound as stimulus poses a challenge both in terms of the low latency needed for the processing not to feel too disturbing and in the lack of a precise control on the nature of the audio presented. While there is a general tendency to move towards more naturalistic stimuli [6][7][8][5], they are still synthetic and not coherent with the real-life environnement.

Furthermore, real-life measurements are challenged by reduced control over environmental conditions, increased motion artifacts, and less control over electromagnetic noise, all of which can adversely affect recording quality.

More recent research on visual stimuli, conducted in more ecological settings [9][10][11][12], have shown that both amplitude and latency of the evoked responses can be modulated by the context in which they are recorded as well as by the degree of ecological validity of the stimulus. Presenting ecologically valid auditory stimuli in real-life settings is however made more difficult by the lack of solutions allowing a controlled stimulation outside of the lab.

In this paper we introduce a new approach which enables research of AEPs in a real-life settings with ecologically valid stimuli. We integrated a portable hearing aid research platform, the Portable Hearing Lab (PHL) [13], and dry-contact ear-EEG [14]. Ear-EEG is a portable and unobtrusive EEG recording technique using dry-contact electrodes embedded in a custom silicone elastomer earpiece [14], which has been shown to be able to record high quality EEG data in a variety of settings [15][16]. Compared to traditional scalp EEG, ear-EEG is less bulky and does not require electrode preparation.

To obtain a portable setup, the ear-EEG earpiece was combined with the PHL behind-the-ear microphones and speakers in a compact solution. In this particular study we aim at showcasing the capabilities of this experimental setup taking as example the recording of the Auditory Steady State Response (ASSR) [17].

Although ASSR has typically been evoked using synthetical stimuli such as amplitude modulated pure tones, amplitude modulated noise [1], or chirps [18]; it has recently been demonstrated that ASSR can be measured in response to amplitude modulated speech signals [19]. Furthermore, it has previously been shown that ASSR can be recorded from ear-EEG [16], and from ear-EEG in real-life settings using amplitude modulated broadband noise [15].

Building on these previous studies, this study advances towards more ecological EEG recordings, by demonstrating that it is possible to elicit and record an ASSR to amplitude modulated ambient sound, with the participant walking around in a shopping center.

There are two primary reasons for selecting the ASSR as the experimental paradigm. First, steady-state responses are characterized by spectral components occurring exclusively at the modulation frequency and its harmonics, facilitating robust and straightforward estimation. Second, ASSRs are widely utilized in physiological hearing assessments as an alternative to behavioral hearing evaluations, in particular when the behavioral testing may be difficult or not possible (e.g. infants or cognitively impaired subjects) thereby enabling hearing assessment without active involvement from the subject [20]. This has the potential to enable unassisted, remote, and longitudinal hearing assessments. When integrated into hearing aid devices, it could facilitate both the initial fitting and ongoing autonomous adjustments in response to progressive hearing loss, reducing the need for in-clinic visits.

## II. METHODS

In the following we describe the details of the hardware platform combining ear-EEG and PHL. To showcase the capabilities of the developed platform we devised an ASSR study, conducted in a busy shopping centre (average SPL 67.9dB, standard deviation 2.6dB).

### A. Hardware Platform

In order to measure ASSR outside of the lab, we integrated the ear-EEG earpieces with the PHL behind-the-ear (BTE) microphone/speaker. To do so we designed and 3D printed (resin printing, Formlabs, tough 1500) a new casing for the BTE headset, able to clip to the ear-EEG earpiece cables. (Figure 1).

**Fig. 1.**
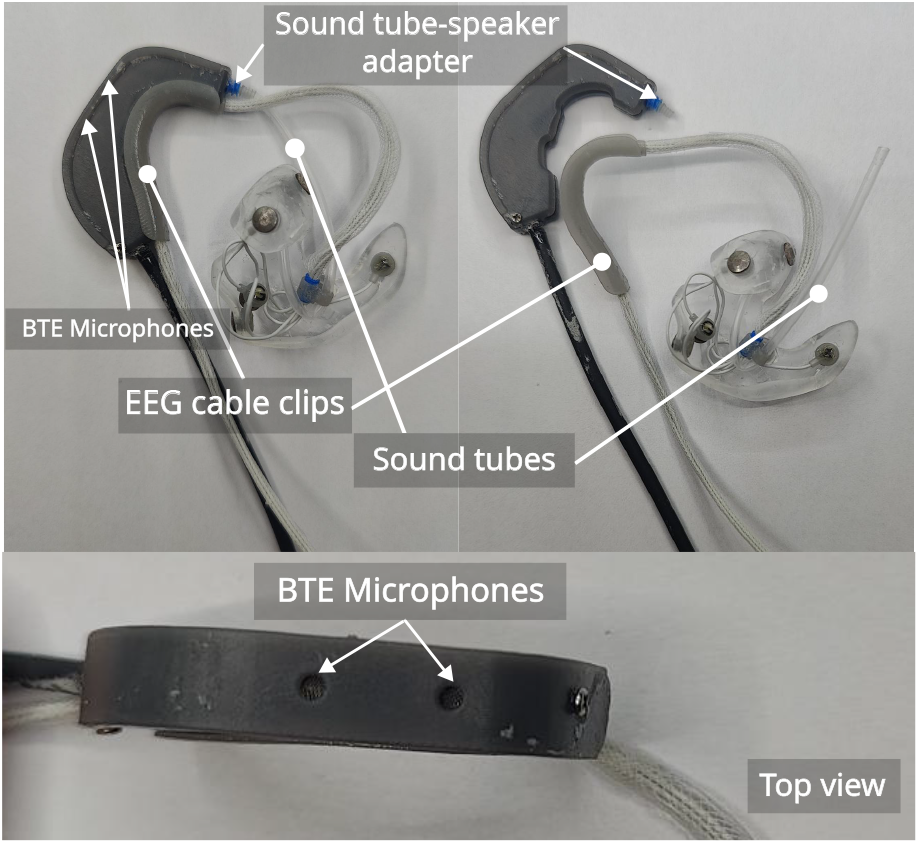
Combined ear-EEG and PHL BTE headset fully assembled and the other taken apart. In the top view it is possible to see the position of the behind-the-ear microphones

In this design, the speaker remains inside the casing and is connected to the earpiece with a soft soundtube (internal diameter 1mm, external diameter 2mm) that joins to a 3D printed adapter inserted in a personalized soft silicone elastomer earpiece thereby strongly attenuating the external sounds.

A soft sound tube is used to decouple the earpiece from the BTE headset as much as possible, minimizing the impact of movements on EEG data quality. A 3D-printed cable relief is inserted into the earpieces to neatly collect and secure the electrode cables and the sound tube. The cables are then routed behind the ear and clipped to the BTE headset to reduce motion related artifacts.

The earpiece is then placed in the participant’s ear, and the modified BTE headset is worn behind the pinna, as shown in Figure 2.

**Fig. 2.**
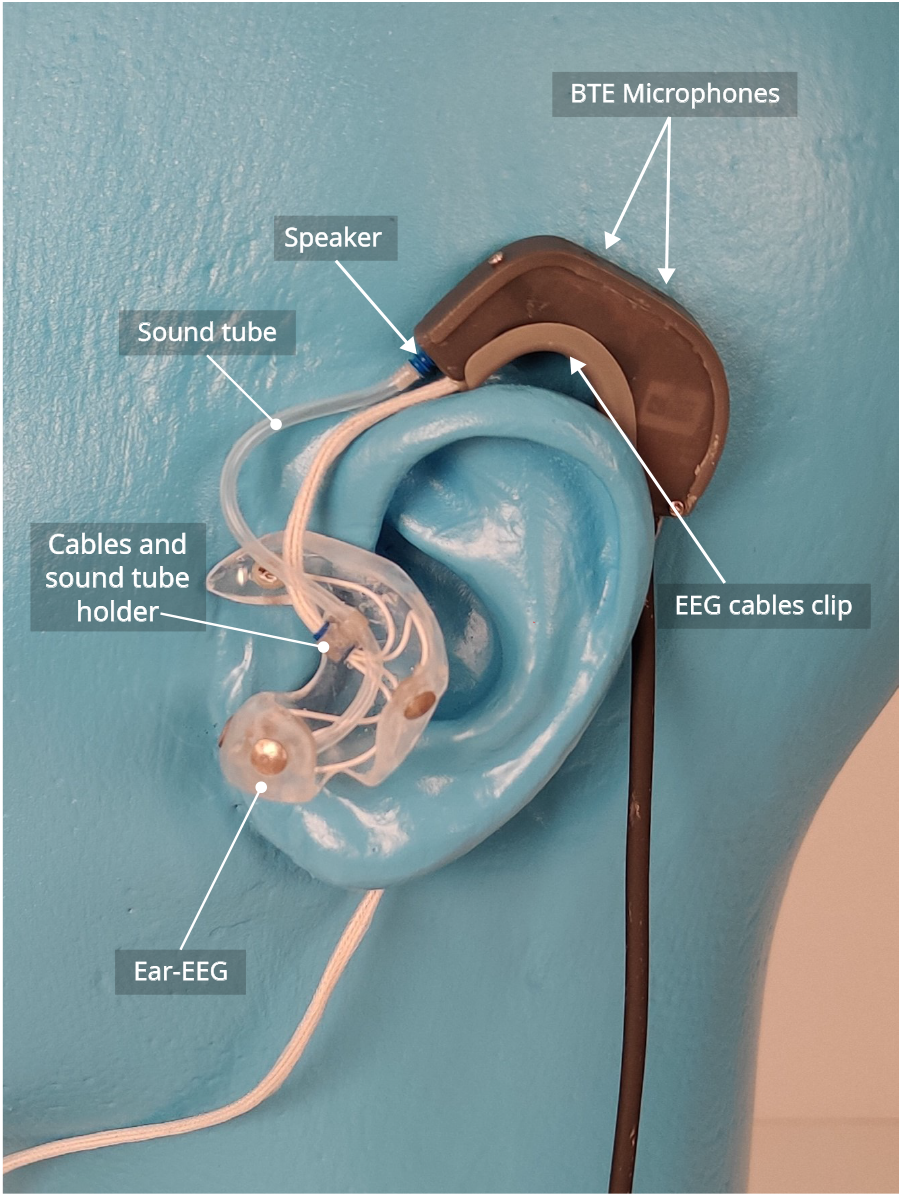
Ear-EEG earpiece with integrated PHL BTE microphone/speaker. Figure 4 shows how it was worn by a participant.

### B. Experimental Setup

The remaining equipment (PHL, EEG amplifier, and trigger box) was carried in a multi-pocket vest worn by the participant.

The EEG was recorded using a 32-channel average reference amplifier (Mobita, TMSi B.V., The Netherlands) at a sampling rate of 1000 Hz. Eight 4 mm Ag/AgCl dry-contact electrodes [21] were placed in the ears (four in the left ear and four in the right) at the locations ELA, ELB, ELF, ELK, ERA, ERB, ERF, and ERK, following the notation established in [16]. For comparison, 24 scalp electrodes were positioned according to the international 10-20 system [22] at locations T7, C3, CP5, P3, CP1, Cz, M1, P7, FC6, P4, P8, CP2, CP6, M2, C4, T8, FC1, FC2, Fz, F4, FC5, Fp2, Fp1, and F3, with the ground electrode at CPz. The scalp electrodes were placed in an electrode cap (Easycap Gmbh, Germany) and prepared with electrode gel (Electro-gel, Electro-Cap International Inc., USA). The ASSR stimulus was presented via in-ear speakers connected to the ear-EEG earpiece using sound tubes (as described in subsection II-A) and consisted of a 40Hz, full depth, amplitude modulation of the ambient sound. 5 normal hearing participants (2 men) aged (25 to 40) were recruited^2^.

### C. Experimental Paradigm

#### 1) Audio Processing

The ambient sound was recorded using the microphones integrated into the behind-the-ear device and processed in real time by the PHL via a custom plugin. This plugin applied a 40 Hz sinusoidal amplitude modulation along with a slight amplification of approximately 1.38 dB.

The custom plugin also created a trigger channel, used to send triggers phase-locked to the amplitude modulation, every ten seconds. The triggers was Manchester coded, with the scheme described in [23]

The processed ambient sound was then played back through the in-ear speakers while the trigger signal was sent to the trigger box for digitalization and galvanic separation [23] from the PHL output jack and then to the Mobita amplifier to be recorded together with the EEG data.

Both the modified ambient sound and the trigger signal were stored in memory as a .wav file for future reference.

Figure 3 depicts the data flow of the experimental setup as well as the audio processing performed by the PHL.

**Fig. 3.**
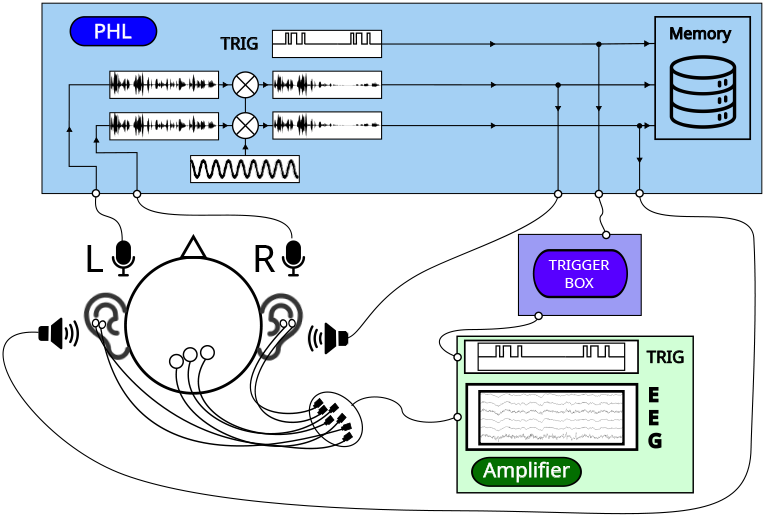
Schematic of the audio processing and data flow between PHL, triggerbox and amplifier in the real-life setup.

#### 2) Tasks

The participants were instructed to perform a series of 3 tasks in the most natural way possible (the only request was to avoid talking if not necessary, to reduce jaw artifacts [24]).

They were asked to:

1. Walk around (20 minutes)
2. Sit on a bench with their eyes open (10 minutes)
3. Sit on a bench with their eyes closed (10 minutes)
4. Sit on a bench with their eyes open (10 minutes)
5. Sit on a bench with their eyes closed (10 minutes)

When walking, they were allowed to enter stores and shop for items (figure 4). While sitting with open eyes, they could use their phone or keep busy as they wished.

**Fig. 4.**
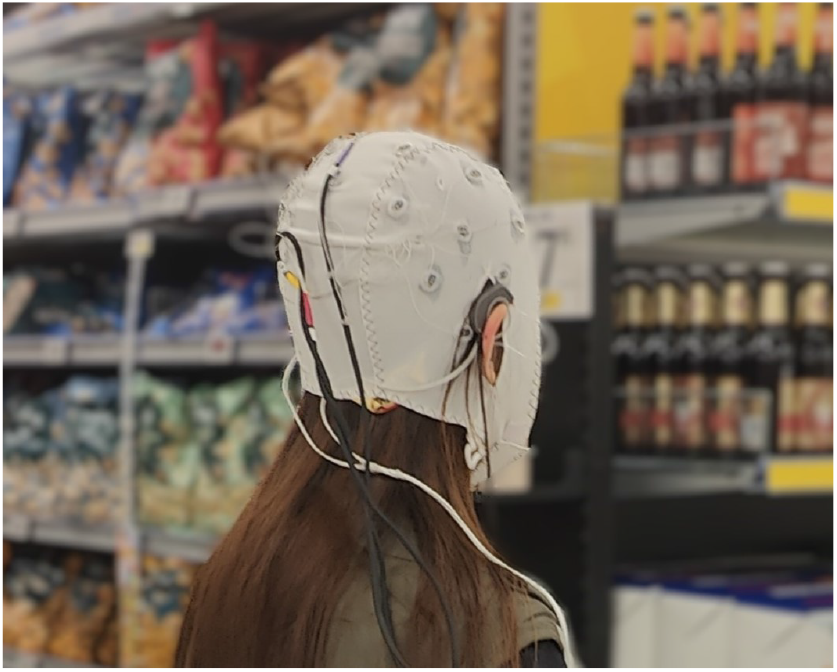
Participant performing the walking task. The subject, depicted in the picture, has given written informed consent to the publication of the image.

The closed eyes task was split in two parts of 10 minutes each, interleave with the open eyes task, to avoid the risk of the participant falling asleep.

### D. EEG Analysis

The collected data were analyzed using the Python MNE package [25][26]. Some channels were saturated due to poor electrode-skin contact and were therefore rejected.

The rest of the channels were combined and referenced based on the following criteria:

- **Scalp**: Average of 8 scalp channels, selected for proximity to the auditory cortex (M1, M2, T7, T8, P7, P8, CP5, CP6), referenced to Fz.
- **Cross-Ear**: Average of the left ear-EEG channels referenced to the average of right ear-EEG channels.
- **In-Ear**: Average spectrum obtained as the average of the spectra for the single channels when referenced to the average of the same ear electrodes, as show in equation 1.

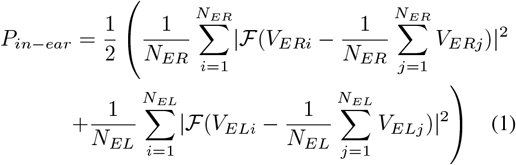

where *N*_*ER*_ and *N*_*EL*_ are the number of electrodes in the right and left ear, respectively (4 for both in this case). *V*_*ERi*_ and *V*_*ELi*_ are the signals from the *i*-th right and left ear-EEG channels respectively and ℱ indicates the discrete Fourier transform.

In all three cases the signal was bandpass filtered between 1Hz and 100Hz using a 4th order Butterworth filter. 5 seconds epochs were extracted using the decoded triggers. All epochs were aligned to phase zero of the amplitude modulation, allowing averaging without signal loss.

Epoch rejection was applied to remove extreme motion artifacts when peak-to-peak amplitude exceeded a threshold of 10mV. The threshold value was chosen expecting a higher noise floor in measurements outside of the lab.

Epochs were averaged, and the Fourier spectrum was computed using pyFFTW [27]. The ASSR response was defined as the power at 40 Hz, while the noise level was calculated as the mean squared amplitude within a ±8 Hz interval around 40 Hz. This interval was selected to provide a sufficiently broad frequency band for noise estimation, while avoiding inclusion of 50 Hz power-line interference and maintaining symmetry around the modulation frequency. The signal-to-noise ratio (SNR) was calculated as the ratio of the ASSR response to the noise level, and its statistical significance was assessed using an F-test.

To determine the needed recording time for a significant response, we repeated the previous analysis for both the walking and the sitting task reducing the number of utilized epochs. This reduction was achieved by picking at random a number of epochs corresponding to the desired recording length, reduced proportionally by the number of bad epochs discarded previously. In this way we accounted for the presence of epochs to be discarded in that time span, assuming them to be homogeneously distributed along the recording.

## III. RESULTS AND DISCUSSION

### A. ASSR while walking in a mall

We report in figures 5a, 5b and 5c the ASSR amplitude (in dB relative to 1nV) for the walking task in the 3 different reference configurations. Each coloured line represents the average over all the non-rejected channels considered in that configuration for a single participant. The thick black line represents the grand average over all participants. Epoch rejection rate was 2.14% for cross-earEEG and 2.05% for in-earEEG, while no epoch was rejected in the scalp configuration.

**Fig. 5.**
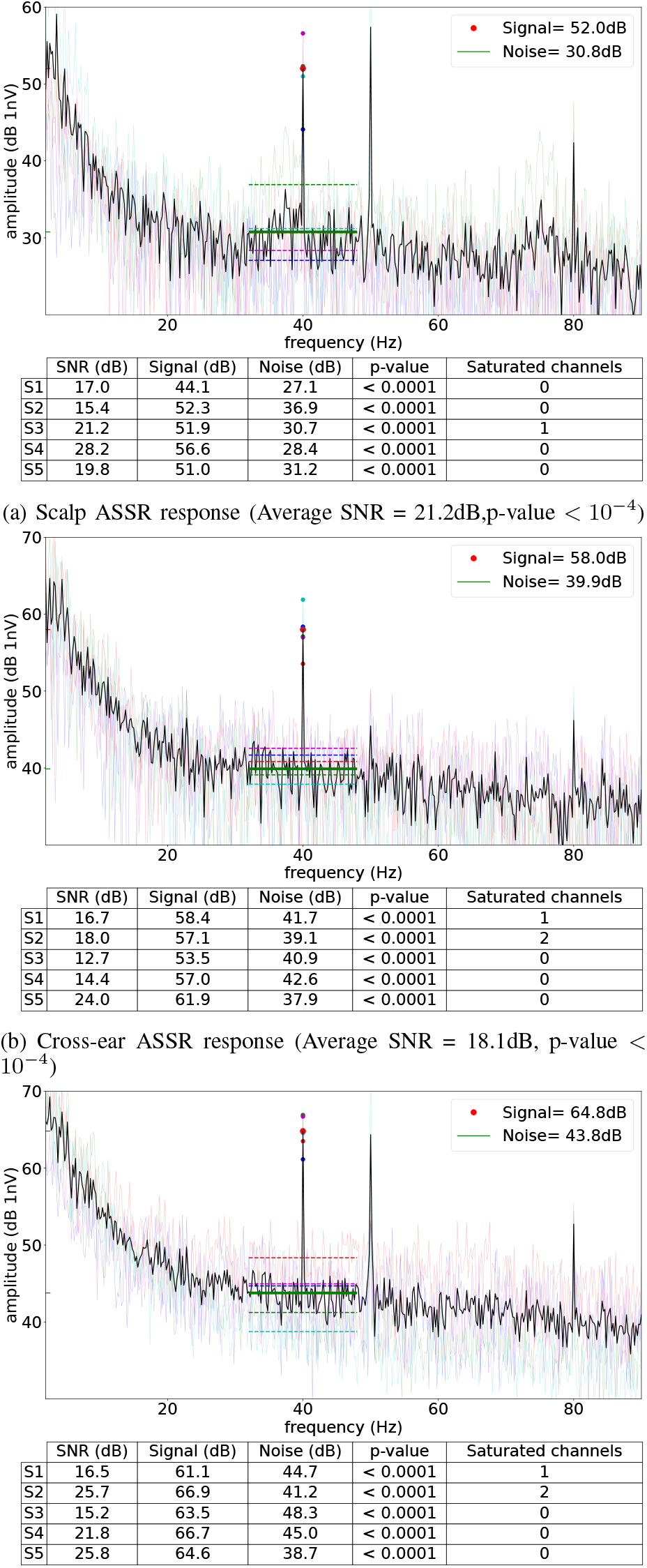
ASSR response in the three different reference configurations for the walking task. The thick black line represents the grand average across participants. Individual participants’ results are shown by the colored lines and in the tables.

The ASSR response is significant (*α* = 0.001) in all three configurations. The other two peaks that can be noticed at 50Hz and 80Hz can be associated respectively with the powerline noise and the second harmonic of the 40Hz ASSR.

The best signal to noise ratio is obtained with the scalp configuration, as expected, followed by the cross-ear and then in-ear. This was to be expected but the fact that the response is still significant for the ear-EEG alone shows how it would be possible to record ASSR data in a real-life scenario, while a participant performs daily life activities, using a couple (or even just one in the case of in-ear referencing) ear-EEG earpieces.

### B. Time for significant results

To explore how long the recording time needs to be to get a significant response in the different electrodes configurations during the 3 different tasks, we performed the previous analysis for subsets of the data with a reduced number of epochs picked as detailed in the methods section. The resulting p-values were marked for 1000 iterations and the time needed to reach a significance level of *α* = 0.01 was taken as the first time point after which the average p-value over the iterations stayed consistently below the threshold.

This times for each participant are reported in figure 6 for the three configurations and the three tasks.

**Fig. 6.**
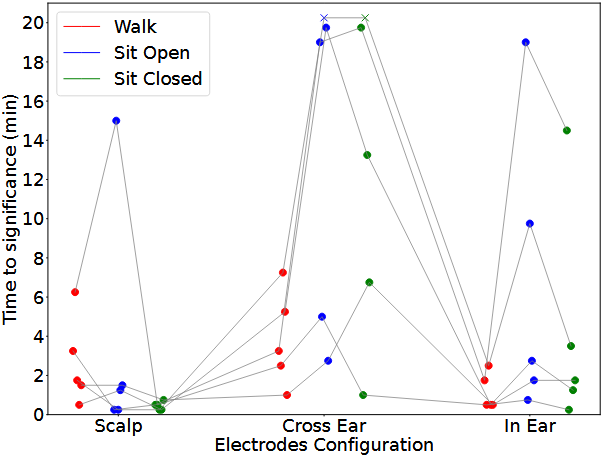
Time needed to reach a significance level of *α* = 0.01 for the three configurations and the three tasks. Each thin grey line represents a single participant. Crosses mark conditions where no significance was reached for the participant after 20 minutes of recording.

As an illustrative example of the procedure used to obtain these times, we report in figure 7 the statistical significance (p-value) vs recording time curves for the three different tasks in the “scalp” configuration. Each point represents one of the random epoch selections, while the curves are the average of the p-values for each participant.

**Fig. 7.**
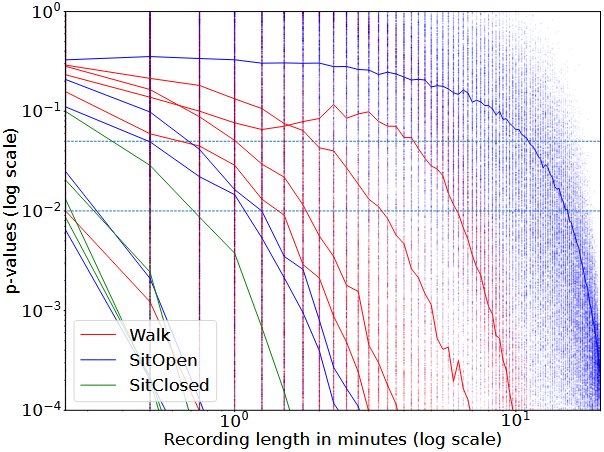
Significance vs recording time curves for the three tasks in the “scalp” configuration. Each point represents one of the permutations, while the curves are the average p-values. The horizontal lines represent the significance levels for *α* = 0.05 and *α* = 0.01.

This showed how a 20 minutes recording time was a strong overestimation for most participants and conditions. When using the scalp configuration, 7 minutes would have been enough to get a p-value below 0.01 for all but one condition in one participant.

For the in-ear configuration, the response was significant with less than 10 minutes of data, for all but two measurements, in both sitting conditions, while it outperformed the scalp configuration in the walking task (4 minutes were enough for all participants), demonstrating the robustness of ear-EEG for real-life measurements.

The cross-ear configuration performed the worst, especially in the sitting tasks. This could be due to the symmetrical topology of the ASSR response [28][29][30], leading to a signal cancellation effect when a cross-head reference is used. However, further measures will be required to verify this hypothesis.

Overall, these results demonstrate that ASSR can be measured using amplitude-modulated ambient sounds in real-life scenarios, within a short recording time, and with a discreet, minimally obtrusive measurement setup

## IV. CONCLUSIONS

In this study, we demonstrated the capabilities of the combined PHL and ear-EEG system to record auditory responses in real-life scenarios.

Specifically, we showed that it is possible to record an ASSR while the subject is walking or sitting outdoors, using amplitude-modulated ambient sound as the stimulus. We found that a recording duration of less than 10 minutes was sufficient to obtain a significant ASSR in the ear-EEG across all three tasks. Furthermore, the results indicated that unconventional recording conditions - such as walking around – required less data to get a significant ASSR response with ear-EEG

The combined PHL and ear-EEG platform opens up the way for future studies aimed at comparing auditory potentials in different conditions, both to get a deeper insight on the brain dynamics in ecological situations and to develop potential integrations between EEG and hearing aid technologies.

## ACKNOWLEDGMENT

This study was supported by the Center for Ear-EEG at Aarhus University

## Footnotes

2 ethical approval number 2024-0660055 from Aarhus University’s Research Ethics Committee (Institutional Review Board)

## Notes

### Competing Interest Statement

The authors have declared no competing interest.

